# BayesAge 2.0: A Maximum Likelihood Algorithm to Predict Transcriptomic Age

**DOI:** 10.1101/2024.09.16.613354

**Authors:** Lajoyce Mboning, Emma K. Costa, Jingxun Chen, Louis-S Bouchard, Matteo Pellegrini

## Abstract

Aging is a complex biological process influenced by various factors, including genetic and environmental influences. In this study, we present BayesAge 2.0, an improved version of our maximum likelihood algorithm designed for predicting transcriptomic age (tAge) from RNA-seq data. Building on the original BayesAge framework, which was developed for epigenetic age prediction, BayesAge 2.0 integrates a Poisson distribution to model count-based gene expression data and employs LOWESS smoothing to capture non-linear gene-age relationships. BayesAge 2.0 provides significant improvements over traditional linear models, such as Elastic Net regression. Specifically, it addresses issues of age bias in predictions, with minimal age-associated bias observed in residuals. Its computational efficiency further distinguishes it from traditional models, as reference construction and cross-validation are completed more quickly compared to Elastic Net regression, which requires extensive hyperparameter tuning. Overall, BayesAge 2.0 represents a notable advance in transcriptomic age prediction, offering a robust, accurate, and efficient tool for aging research and biomarker development.

## 1 Introduction

Aging, once considered an unavoidable process in the cycle of life, is now increasingly viewed as a condition that can be modified. The growing field of aging research has revealed that biological age (BA), which is distinct from chronological age (CA), can be influenced by various genetic, environmental, epigenetic, and lifestyle factors.[1–3] While CA is simply the number of years an individual has lived, BA measures the extent of cellular and molecular aging, providing insights into an individual’s overall health, aging process, and potential lifespan.[4, 5]

In the ‘omics’ era, various epigenetic, transcriptomic, metabolomic, proteomic, microbiomic, and glycomic clocks have been developed to identify biomarkers of aging.[6] Understanding the mechanisms of aging and developing reliable biomarkers for age prediction are crucial for advancing biomedical research by enabling the identification of key molecular and cellular pathways involved in the aging process.[7] These advances facilitate the development of targeted therapies and interventions, support the early detection of age-related diseases, and help optimize personalized medicine approaches aimed at improving healthspan and reducing the burden of agerelated conditions.[8] Moreover, these biomarkers provide a critical tool for evaluating the effectiveness of interventions designed to modify biological age, allowing for the assessment of therapies intended to slow or reverse aspects of the aging process.[9, 10]

Transcriptomic age (tAge), an estimate of biological age based on gene expression profiles, has emerged as a promising biomarker, reflecting the molecular changes that accompany aging.[11, 12] Previous transcriptomic clocks based on neural networks require large datasets to train accurate models, which can be problematic as access to such data is often limited.[13, 14] This limitation can hinder the development and validation of robust models, potentially affecting their accuracy and generalizability in real-world applications. As an alternative, Elastic Net regression, a hybrid model combining Ridge and LASSO regression, is often used.[3, 15, 16] This method adds a single penalty term controlled by a parameter *α* to the ordinary least squares (OLS) objective function, where the penalty is a combination of the L1 (LASSO) and L2 (Ridge) norms. Additionally, Elastic Net includes an L2 ratio parameter, which determines the strength of the Ridge penalty relative to the overall regularization applied to the model. However, although the aging clocks based on Elastic Net regression can predict chronological age accurately, the assumption of linearity in methylation or transcriptomic patterns with age may not fully capture the underlying biological processes. These models also tend to overfit, lack interpretability, and require extensive hyperparameter tuning. Additionally, they are sensitive to missing data, making age prediction challenging when features in the testing sample are absent in the trained model. In bulk RNA sequencing, gene expression data are typically represented as count data, which are discrete and non-negative. To appropriately model such data, statistical frameworks often employ Poisson or negative binomial distributions, which account for the inherent variability in gene expression levels across different samples.[17–22] The Poisson distribution, in particular, is commonly used when the mean and variance of the count data are approximately equal, making it a suitable choice for many gene expression studies.

To address the shortcomings of linear regression models used in epigenetic clocks, we previously developed BayesAge, a maximum likelihood algorithm to predict epigenetic age.[23] BayesAge was inspired by the scAge methodology, which was originally designed for single-cell DNA methylation analysis.[24] BayesAge utilized locally weighted scatterplot smoothing (LOWESS) to capture non-linear methylation-age dynamics and a binomial distribution to model bisulfite sequencing count data.

In this work, as a proof of principle, we extend the BayesAge framework to predict transcriptomic age. Building on the foundations of its predecessor, BayesAge 2.0 leverages a robust statistical framework to model the relationship between gene expression and age. By incorporating prior biological knowledge, our algorithm provides reliable and biologically meaningful age predictions.

The primary contributions of BayesAge 2.0 are threefold. First, it employs a maximum likelihood approach to estimate model parameters, ensuring an optimal fit to the observed data. Second, it models gene expression counts using a Poisson distribution and incorporates LOWESS to capture the non-linear relationships between gene expression levels and age. Finally, BayesAge 2.0 can predict biological age even when some features or measurements are missing from the dataset.

The dataset used in this study was obtained from the Tabula Muris Consortium [25, 26]. This dataset contains bulk RNA sequencing of 17 organs from C57BL/6JN mice across the organism’s lifespan. Tissues were collected from both male and female mice. For males, samples were collected from 4 mice at each of the following ages: 1, 3, 6, 9, 12, 15, 18, 21, 24, and 27 months. For females, samples were collected from 2 mice at each of the following ages: 1, 3, 6, 9, 12, 15, 18, and 21 months. This dataset encompasses stages from early development at 1 month old to maturity at 3–6 months, and extends through aging up to the median lifespan of 27 months. The organs sampled from each mouse included bone, brain, brown adipose tissue (BAT), gonadal adipose tissue (GAT), heart, kidney, limb muscle, liver, lung, bone marrow, mesenteric adipose tissue (MAT), pancreas, skin, small intestine, spleen, subcutaneous adipose tissue (SCAT), and white blood cells (WBC).

## 2 Methods

### 2.1 BayesAge transcriptomic clock algorithm overview

As in our previous version, we re-implemented the BayesAge framework for transcriptomic age prediction in two steps: training and prediction, as illustrated in Figure 1.

**Fig. 1.**
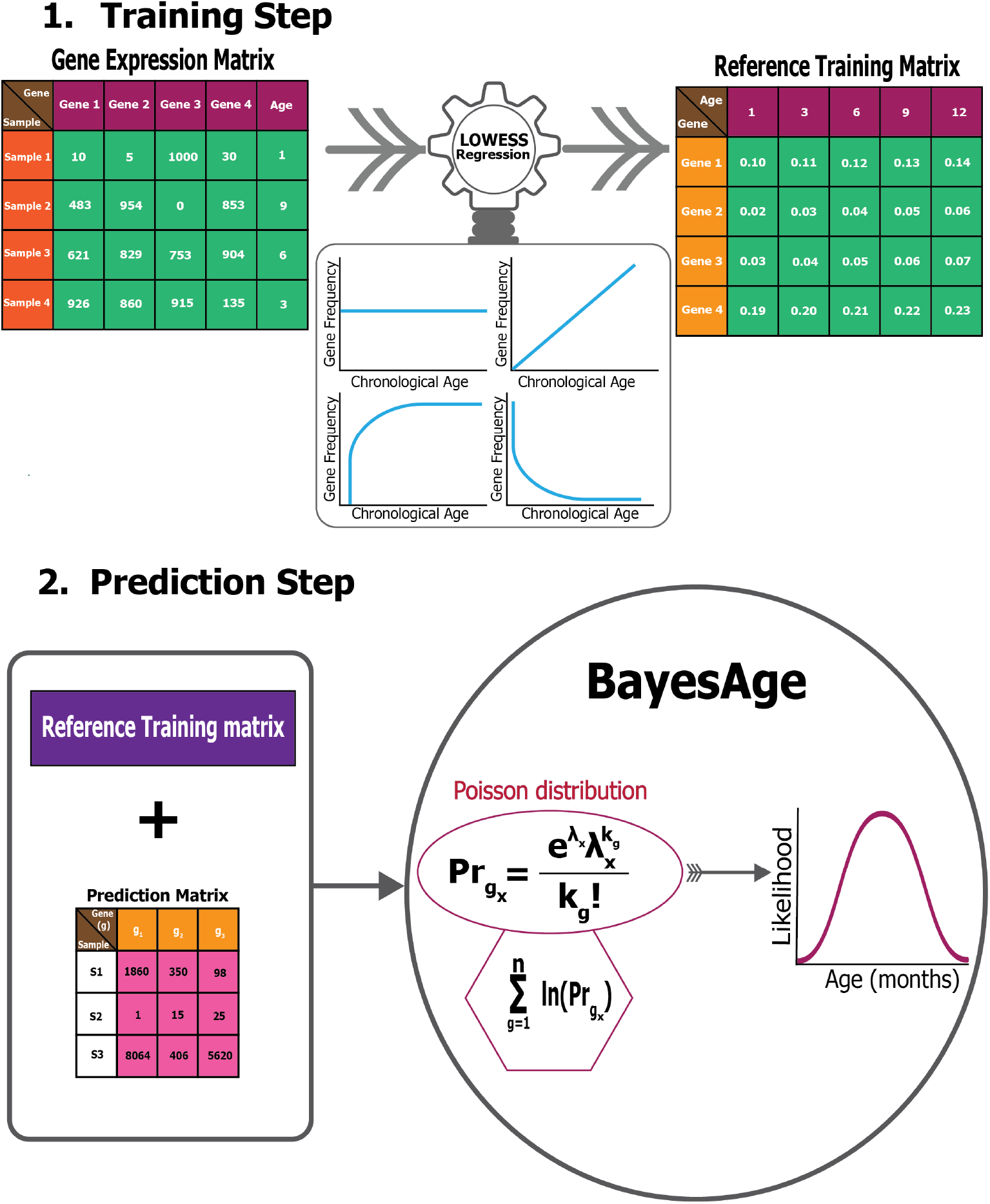
BayesAge transcriptomic clock framework.

In the training step, we first normalize the raw gene expression counts using frequency count normalization (Eq. 1). Frequency count normalization adjusts the raw counts of features, such as genes, so that they are comparable across samples. This is done by dividing each count by the total number of counts in the sample, converting raw counts into relative frequencies or proportions.

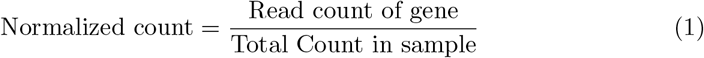

Next, we used LOWESS regression to fit the trend between gene frequency levels for each gene and age, accounting for the non-linearity of the frequency patterns. The *τ* parameter determines the smoothness of the LOWESS fit. **statsmodels.api**’s LOWESS function was used in the implementation of BayesAge. After computing the fit for each gene, we calculate the correlation between frequency levels and age using the Spearman rank correlation, which is robust to non-linear trends. We select the top genes to include in the prediction step using the absolute value of the correlation. At the end of this process, the training model consists of N genes, and their frequency levels across ages, ranging from 1 to 27 months in increments of 3 months, based on a predetermined *τ* parameter (0.7 in this study) for the LOWESS fit. The value of the *τ* parameter is empirically determined such that we avoid over fitting when *τ* is closer to zero and under fitting when *τ* is closer to 1.

In the prediction step, the reference matrix is intersected with the genes expressed in a specific sample. The raw counts from the gene expression matrix produced for any downstream Bulk RNA-seq data analysis can be used for the prediction. For the chosen age-associated genes, we propose that the chance of detecting the observed raw counts, given the intended frequency level for a specific age based on the trained model, follows a Poisson distribution.[17–19, 21, 22] To compute the probability of observing the counts measured across all the genes identified from the intersection with the training matrix, we compute the product of these probabilities. To prevent underflow errors during computation, a sum of logarithms replaces the product of individual genes, yielding a singular probability of each age.

Utilizing these pre-identified, ranked age-associated genes, the framework calculates the likelihood of observing each age in a single subject, spanning an age spectrum of 1-27 months, at 3-month intervals. As such, for each subject, we compute an age-likelihood distribution, with the maximum likelihood age interpreted as the transcriptomic age for that subject. In this framework, Pr_*gene*_ represents the gene expression probability for a distinct gene g at a specific age, aggregated from 1 gene to N total genes. The associated probability for a unique gene state is:

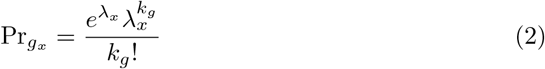

where:

*x*: specific age in the age likelihood probability distribution.

*λ*_*x*_: expected count at age *x*.

*k*_*g*_: observed count that was measured for the test sample *ϕ*.

### 2.2 Validation

For both BayesAge and Elastic Net regression, we implemented Leave One Out Cross Validation (LOOCV) to validate the age predictions.

For the age prediction using Elastic Net regression, we first log-transformed the data using **numpy’s** *log1p* function to stabilize the variance. Subsequently, we scaled the features of the gene expression matrix using the **StandardScaler** function from the **sklearn.preprocessing** package. The standard scaler ensures that the features are standardized to have a mean of zero and a standard deviation of 1. We also used the **LeaveOneOut** function from the **sklearn.model selection** package to implement the cross-validation. Since Elastic Net regression requires hyperparameter tuning, we implemented a parameter grid search for the alpha parameter (which controls the overall strength of the regularization to the loss function, combining both L1 and L2 penalties) and the l1 ratio (which determines the relative contribution of L1 versus L2 regularization). This search was conducted in two passes. In the first pass, to find the best parameters, the alpha parameter values tested ranged from [1e-5, 1e-4, 1e-3, 1e-2, 1e-1, 1.0, 10.0, 100.0], and the l1 ratio values tested ranged from 0 to 1 in steps of 0.1. The maximum number of iterations was set to 10,000. Additionally, the scoring function was the mean absolute error (MAE). Once the best parameters were identified, in the second pass, we increased the maximum number of iterations to 100,000 to ensure the objective function converged and to make the final age predictions.

### 2.3 Software

In BayesAge 1.0, we implemented three main functions to predict the epigenetic age of organisms: **construct_reference, load_cgmap_file**, and **bdAge**. Here, we have updated the main functions as follows: To predict the epigenetic age, one should use the **epigenome_reference** to construct the reference matrix using bulk methylation data in conjunction with the **bdAge** function. To predict the transcriptomic age, one should use the **transcriptome_reference** along with the **pdAge** function.

## 3 Results

### 3.1 BayesAge Framework

We extended our previous model, BayesAge, originally designed to predict epigenetic age, to predict transcriptomic age. Our adapted BayesAge for tAge has two steps. In the first step, we train the model using raw gene expression counts, and in the second step, we predict the age of a sample. In the training step, we select genes in the mouse transcriptome that have age-associated patterns. As shown in Figure 2, the gene expression patterns in the mouse brain are non-linear; therefore, using a linear function to describe these changes would not accurately capture them, justifying the need for non-linear functions to model these dynamic changes. To identify the most significantly age-associated genes, we used the Spearman rank correlation, which is a non-parametric method that does not assume linearity. In the mouse brain, we observed that for many of these genes, the association of gene frequency with age is non-linear, with rates of change that increase with age. In general, for the brain we identified 16,108 genes with a positive Spearman rank, 7,874 genes with a negative Spearman rank, 3 genes with a zero Spearman rank, and 631 genes with a correlation coefficient not defined due to the input array or gene level being constant in the brain.

**Fig. 2.**
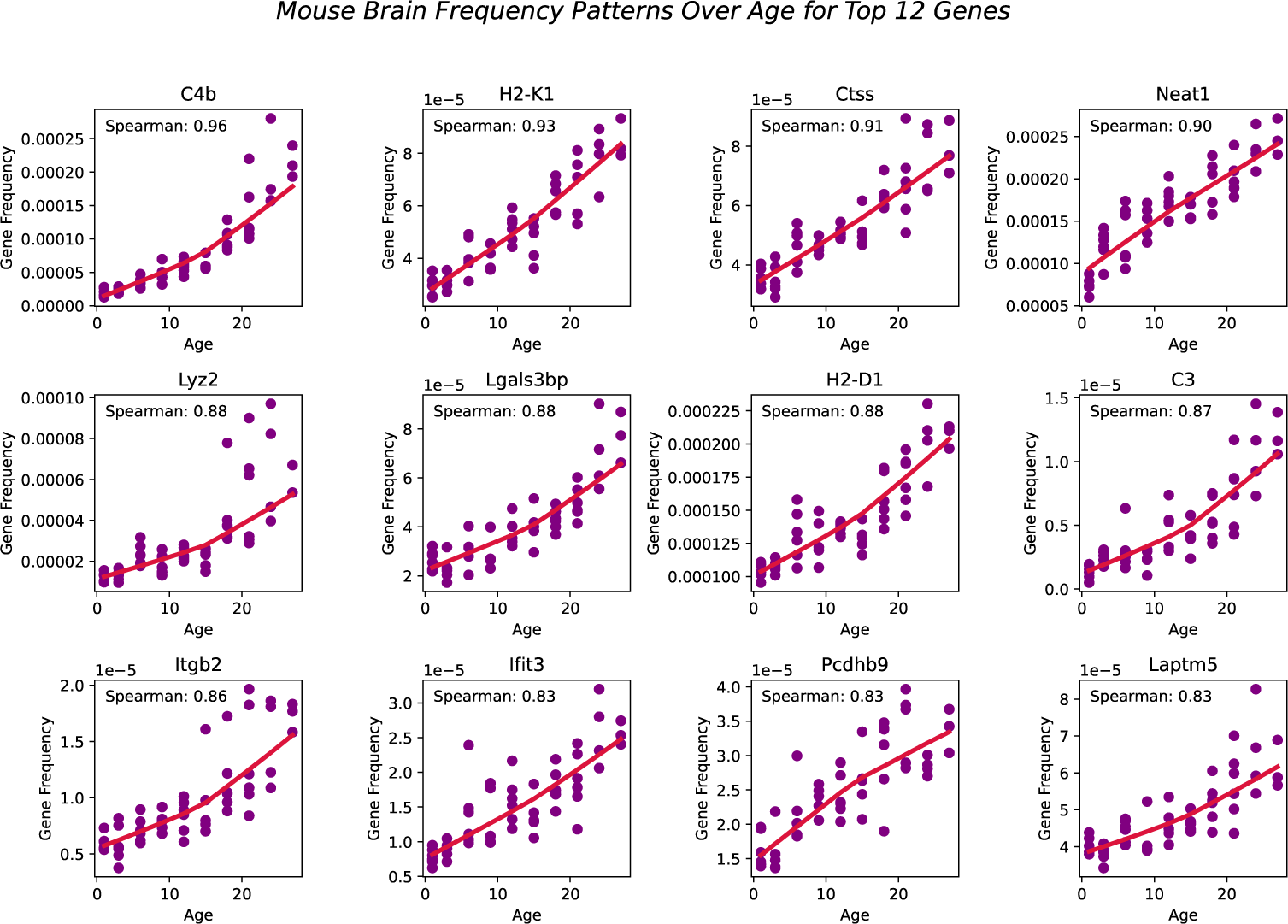
Top 12 Genes fitted with Spearman correlation values using LOWESS regression (red line) with a tau parameter value of 0.7.

BayesAge, recognizing these non-linear trends, uses LOWESS regression to model the trend lines. This method utilizes locally weighted linear regression to estimate a smoothed line through the data. The extent of this smoothing is governed by the tau parameter, which determines the size of the local neighborhood window for each local linear regression fit. We empirically tested different tau values and visually inspected the model fit. A tau value of 0.7 was chosen, allowing the fit to adaptively capture the nonlinear gene frequency pattern variations across the age spectrum without succumbing to overfitting. The trend lines of these fits represent our aging model, or the expected frequency with age at each of these genes.

Our prediction step allows us to estimate the age of a sample using the training model. This approach builds on the count-based nature of gene expression data to estimate age from the gene frequency data of a single sample. We propose a Bayesian framework for estimating the most likely age of an individual by computing the probability of the observed counts of a gene for any given age and selecting the age that maximizes this probability.

To estimate the probability of the observed counts based on the expected gene frequency levels of a single gene at a specific age, we use a Poisson distribution. This choice is rooted in the nature of RNA-seq data, where count data corresponds to the number of reads mapped to each gene. Each read represents an independent observation, and the number of reads observed for a particular gene can be modeled as a Poisson process.[17–19, 21, 22] The expected count for each gene, which serves as the parameter for the Poisson distribution, is proportional to the gene’s true expression level and is derived from the gene frequencies observed in the sample. Thus, the observed counts across genes follow a Poisson distribution, where the rate parameter reflects the expected counts based on gene frequencies.

To estimate the probability of a specific age for a sample, we computed the likelihood of our observed counts across multiple genes for any given age by taking the product of the probabilities of each gene. In practice, this product is computed by summing the logarithms of the probabilities. Finally, we identified the age that maximizes this probability.

We found that the transcriptomic clock from the mouse brain was the best performing clock with our model, achieving an R-squared value of 0.908 and an MAE of 1.83 months, as shown in Figure 3. Surprisingly, the accuracy of predictions for the brain remains stable as the number of top correlated genes used for prediction increases; even when using the top 1000 genes, we still obtained an R-squared value of 0.860.

**Fig. 3.**
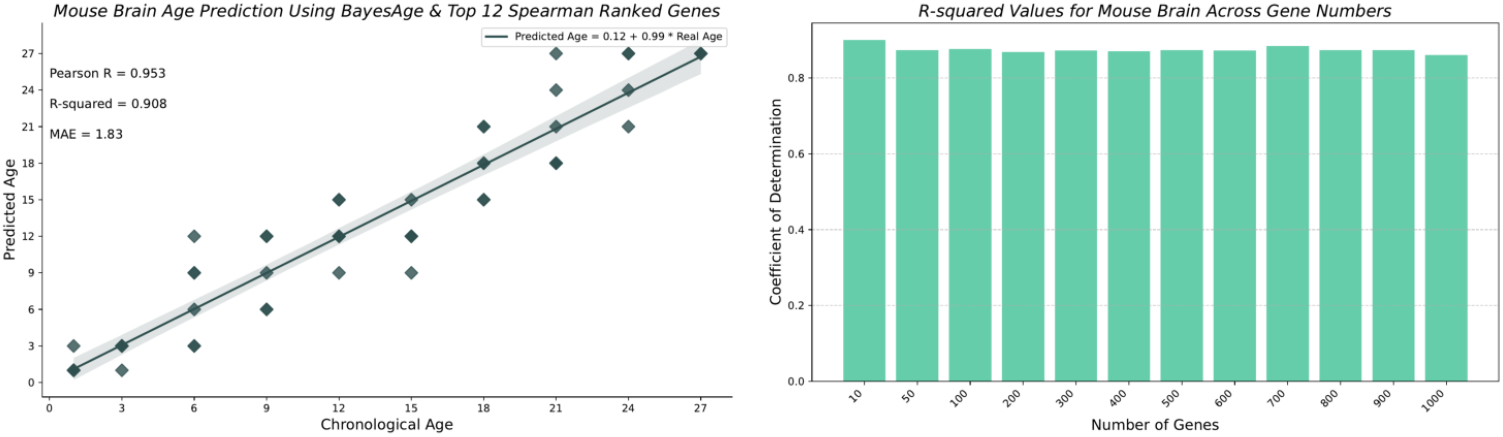
Mouse Brain age predictions using BayesAge along with the Pearson r, R-squared and MAE values in months.

Figure 4 highlights the performance of the best-performing clocks, along with the number of genes used across all the tissues. The worst-performing clock was for the skin tissue, with an R-squared of 0.265 and an MAE of 5.47 months. The number of genes tested ranged from the top 12, 15, 20, 25, and 30 for all the tissues. From the top correlated genes used for age prediction using BayesAge, we identified only two genes that had a positive correlation across all the tissues: *Mgmt* (ENSMUSG00000054612) and *Bhlhe40* (ENSMUSG00000030103). We also identified only one gene that had a negative correlation across all the tissues: *Col1a1* (ENSMUSG00000001506), as shown in the excel file with spearman ranked results used in this study.

**Fig. 4.**
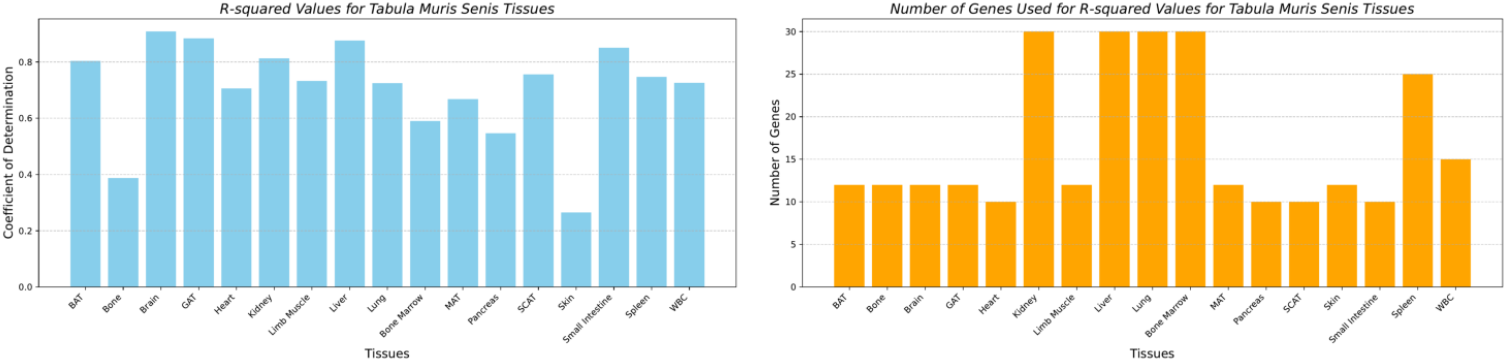
R-squared values of the best performing clocks along the number of genes used across tissues of the Tabula Muris Senis.

### 3.2 Comparison of BayesAge with Elastic Net Regression

We used the best-performing transcriptomic clock developed with BayesAge and implemented the same transcriptomic clock using Elastic Net to evaluate the performance of our model. As shown in Figure 5, LOOCV coupled with MAE was employed to validate the outcomes of both models. The results show that BayesAge has a higher coefficient of determination (R-squared) compared to the Elastic Net model using only the top 12 Spearman-ranked genes. Interestingly, BayesAge outperforms the Elastic Net model for the brain tissue, with an R-squared value of 0.908 and an MAE of 1.83 months, compared to an R-squared value of 0.892 and an MAE of 2.190 for the Elastic Net model.

**Fig. 5.**
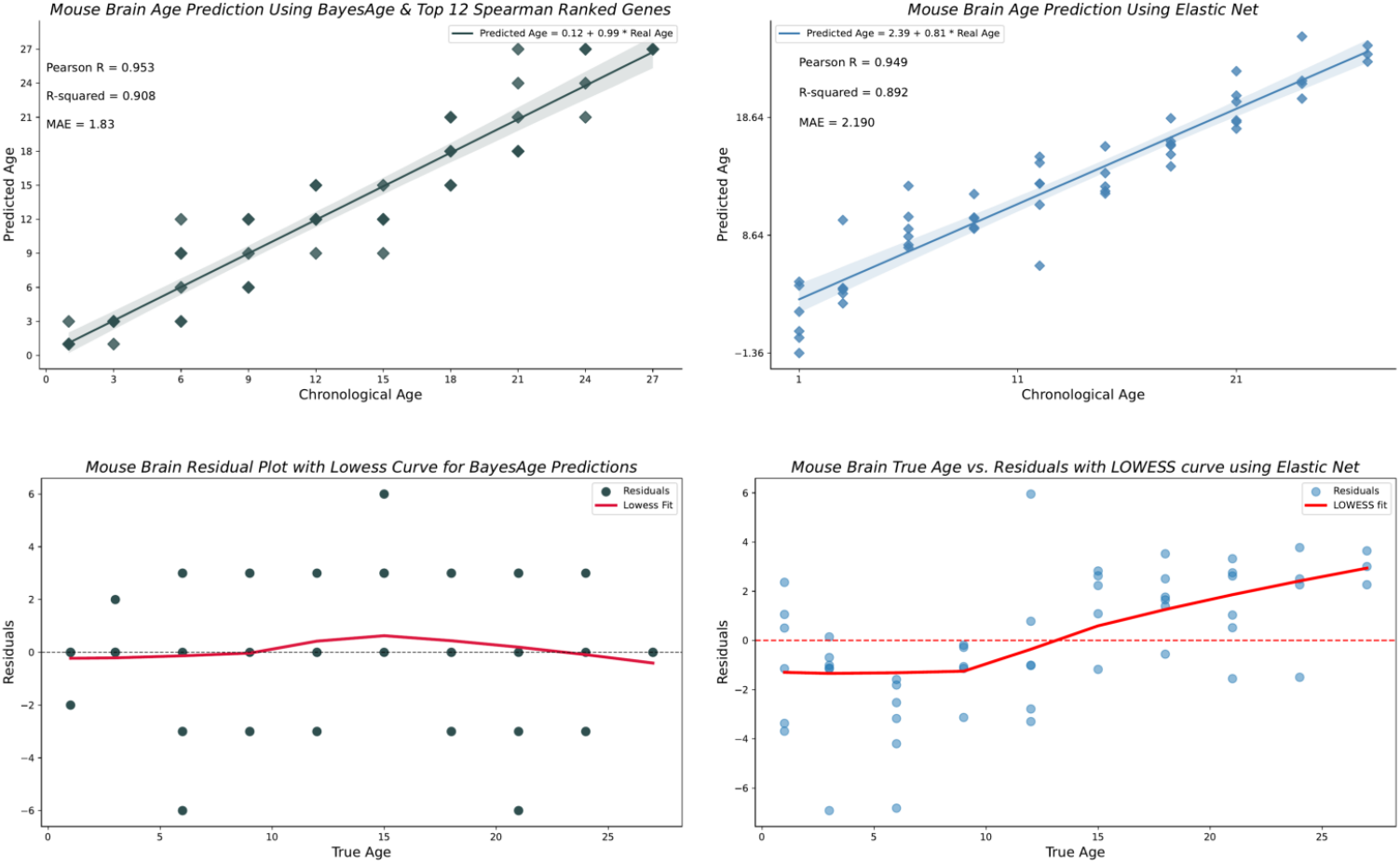
Comparison of BayesAge and Elastic Net age predictions for the Brain tissue.

Age bias in the predictions is another crucial metric. Age acceleration is usually defined as the difference between the biological age determined by the machine learning model and the chronological age, with these differences often associated with agerelated diseases. However, the residuals should not be correlated with age. Therefore, we evaluated the residuals of the brain’s Elastic Net model and BayesAge model to identify any age-related biases. As shown in Figure 5, the LOWESS fit of BayesAge, with a tau parameter of 0.7, shows low age-associated biases. In contrast, the Elastic Net model residuals exhibit an age-residual pattern resembling a logistic function. This absence of quantitative age-associated residual patterns is another advantage of the non-linear BayesAge over Elastic Net regression and other similar linear models. Table 1 shows the overall performance of BayesAge compared to Elastic Net.

**Table 1.** Comparison of BayesAge results vs. Elastic Net for all the Tabula Muris Senis tissues.

| Tissue | BayesAge |  |  |  | Elastic Net |  |  |
| --- | --- | --- | --- | --- | --- | --- | --- |
|  | # of genes used | Pearson R | R-squared | MAE | Pearson R | R-squared | MAE |
| BAT | 12 | 0.897 | 0.804 | 2.30 | 0.927 | 0.841 | 2.489 |
| Bone | 12 | 0.623 | 0.388 | 5.52 | 0.836 | 0.637 | 3.899 |
| Brain | 12 | 0.953 | 0.908 | 1.83 | 0.949 | 0.892 | 2.190 |
| GAT | 12 | 0.941 | 0.884 | 1.87 | 0.955 | 0.906 | 2.034 |
| Heart | 10 | 0.840 | 0.706 | 2.17 | 0.578 | 0.267 | 4.829 |
| Kidney | 30 | 0.902 | 0.813 | 2.73 | 0.947 | 0.878 | 2.179 |
| Limb Muscle | 12 | 0.856 | 0.733 | 2.96 | 0.930 | 0.850 | 2.468 |
| Liver | 30 | 0.936 | 0.876 | 1.98 | 0.944 | 0.872 | 2.265 |
| Lung | 30 | 0.851 | 0.725 | 2.93 | 0.939 | 0.872 | 2.265 |
| Bone Marrow | 30 | 0.768 | 0.590 | 3.71 | 0.836 | 0.637 | 3.899 |
| MAT | 12 | 0.817 | 0.668 | 3.33 | 0.922 | 0.814 | 2.964 |
| Pancreas | 10 | 0.740 | 0.547 | 3.67 | 0.893 | 0.734 | 3.398 |
| SCAT | 10 | 0.870 | 0.756 | 2.73 | 0.883 | 0.774 | 3.032 |
| Skin | 12 | 0.515 | 0.265 | 5.47 | 0.894 | 0.780 | 2.965 |
| Small Intestine | 10 | 0.922 | 0.851 | 1.84 | 0.944 | 0.880 | 1.899 |
| Spleen | 25 | 0.864 | 0.747 | 3.07 | 0.920 | 0.826 | 2.564 |
| WBC | 15 | 0.852 | 0.726 | 2.67 | 0.923 | 0.829 | 2.496 |

### 3.3 Computational Performance

On the computational front, the time required to identify the optimal hyperparameters for the age prediction using Elastic Net is over 90 minutes for LOOCV. In contrast, the time to construct one reference using BayesAge’s **transcriptome reference** is about 30 seconds, totaling approximately 27 minutes to implement a LOOCV strategy using BayesAge for the over 50 samples in the Tabula Muris Senis dataset per tissue. With no need to optimize hyperparameters other than explicitly selecting the number of Spearman-correlated genes to use in the tAge prediction, and offering better interpretability and less age bias in age prediction, BayesAge demonstrates a clear computational advantage over Elastic Net.

## 4 Discussion

In this study, we introduced BayesAge 2.0, an enhanced version of our maximum likelihood algorithm designed for predicting tAge from gene expression data. This advancement builds upon the original BayesAge framework, initially developed for epigenetic age prediction, by incorporating a Poisson distribution to model count data and utilizing LOWESS smoothing to address non-linear gene-age relationships.

BayesAge offers several notable advantages over traditional linear models, such as Elastic Net regression. First, our method employs a maximum likelihood approach, ensuring a robust fit to the observed gene expression data. By using a Poisson distribution, BayesAge effectively captures the count-based nature of RNA-seq data, which aligns with the discrete nature of gene expression measurements. This approach enhances the accuracy of age predictions by directly addressing the probabilistic characteristics of gene counts. Additionally, age prediction in the absence of features or genes is now possible with our approach.

The application of LOWESS smoothing enables BayesAge to capture non-linear trends in gene expression data, which is essential given the complexity of aging changes. As evidenced by our results, the gene expression patterns exhibit non-linear relationships with age, particularly in tissues such as the brain. By fitting these patterns with LOWESS, our model adapts to variations in gene frequency, offering more accurate age estimates compared to linear regression models that may oversimplify these relationships.

Our comparison of BayesAge with Elastic Net regression highlights its superior performance for specific tissues. For the brain tissue, BayesAge achieved an R-squared value of 0.908 and a mean absolute error (MAE) of 1.83 months, outperforming the Elastic Net model, which had an R-squared of 0.892 and an MAE of 2.190 months. This improvement underscores BayesAge’s ability to capture complex age-related changes more effectively for specific tissues.

Furthermore, BayesAge 2.0 addresses issues related to age bias in predictions. The residuals from our model exhibit minimal age-associated bias, in contrast to the more pronounced age bias observed in the Elastic Net model. The absence of residual bias indicates that the model’s predictions are not systematically overestimating or underestimating the actual values across different ranges of the data. This is important, as we interpret the residual as a biological process such as age acceleration or deceleration, but if the age prediction is biased, the biological process inferred from the model will also be biased. This suggests that on average, the model’s predictions are accurate and don’t favor higher or lower estimates. Additionally, an unbiased model is more likely to generalize well to new, unseen data. This is due to the fact that a model with residual bias may perform well on the training data but poorly on new data because it has learned to overestimate or underestimate in a particular matter. This reduced bias indicates that BayesAge 2.0 provides more interpretable and reliable age estimates that is more generalizable to unseen data, which is crucial for applications in aging research and clinical settings.

BayesAge also demonstrates significant computational efficiency. While Elastic Net regression involves extensive hyperparameter tuning and can be time-consuming, BayesAge’s approach is more streamlined. Constructing a reference matrix and performing cross-validation with BayesAge is substantially faster, with a total time of approximately 27 minutes for the Tabula Muris Senis dataset for a LOOCV.

The interest in aging clocks stems from their potential use as biomarkers in intervention studies aimed at extending lifespan. Future studies may explore the use of BayesAge for lifespan intervention studies using dietary changes, pharmacological treatments, genetic modifications, or lifestyle alterations in aging organisms such as mice, fishes and humans.[12, 27–30].

In summary, BayesAge represents a notable advancement in transcriptomic age prediction, offering improved accuracy, reduced bias, and greater computational efficiency compared to traditional linear models. By leveraging a maximum likelihood approach, a Poisson distribution for count data, and LOWESS smoothing for non-linear trends, our model provides a comprehensive tool for understanding and predicting biological age at the transcriptomic level. As the field of aging research continues to evolve, particularly within the genetic aging community, BayesAge is well-positioned to offer valuable insights to support the development of more effective therapies for age-related studies.

## Declarations

## 4.1 Acknowledgements

Lajoyce Mboning is a Eugene V. Cota-Robles, Warren Alpert Computational Biology and AI Network and a NSF NRT fellow. This work was supported by the National Institutes of Health (NIH) Training Grant in Genomic Analysis and Interpretation T32HG002536.

## 4.3 Data availability

The data used in this study is deposited in the Gene Expression Omnibus (GEO) database, under accession number GSE132040. A subset of the raw counts used in this study is available on the official GitHub repository.

## 4.4 Code availability

The code for this study is available at: https://github.com/lajoycemboning/BayesAge2.0

## 4.5 Author contribution

**LM**: Conceptualization, Data curation, Formal Analysis, Investigation, Methodology, Software, Validation, Visualization, Writing–original draft, Writing–review and editing. **EKC**: Data curation, Writing–review and editing. **JC**: Writing–review and editing. **LB**: Supervision, Writing–review and editing. **MP**: Conceptualization, Methodology, Supervision, Writing–review and editing.

## References

[1] Mitchell, W., Goeminne, L.J., Tyshkovskiy, A., Zhang, S., Chen, J.Y., Paulo, J.A., Pierce, K.A., Choy, A.H., Clish, C.B., Gygi, S.P., Gladyshev, V.N.: Multiomics characterization of partial chemical reprogramming reveals evidence of cell rejuvenation. eLife 12, 90579 (2024) 10.7554/eLife.90579.3

[2] Lu, Y., Tian, X., Sinclair, D.: The information theory of aging. Nat Aging 3(12), 1486–1499 (2023) 10.1038/s43587-023-00527-6. Epub 2023 Dec 15

[3] Peters, M., Joehanes, R., Pilling, L., et al.: The transcriptional landscape of age in human peripheral blood. Nature Communications 6, 8570 (2015) 10.1038/ncomms9570

[4] Bafei, S.E.C., Shen, C.: Biomarkers selection and mathematical modeling in biological age estimation. npj Aging 9(13) (2023) 10.1038/s41514-023-00110-8

[5] Sagers, L., Melas-Kyriazi, L., Patel, C.J., Manrai, A.K.: Prediction of chronological and biological age from laboratory data. Aging (Albany NY) 12(9), 7626–7638 (2020) 10.18632/aging.102900

[6] Rutledge, J., Oh, H., Wyss-Coray, T.: Measuring biological age using omics data. Nature Reviews Genetics 23, 715–727 (2022) 10.1038/s41576-022-00511-7

[7] Guo, J., Huang, X., Dou, L., et al.: Aging and aging-related diseases: from molecular mechanisms to interventions and treatments. Sig Transduct Target Ther 7(391) (2022) 10.1038/s41392-022-01251-0

[8] Chen, R., et al.: Biomarkers of ageing: Current state-of-art, challenges, and opportunities. MedComm – Future Medicine 2(2), 1–12 (2023) 10.1002/mef2.50

[9] Moqri, M., Herzog, C., Poganik, J.R., Aging Consortium, B., Justice, J., Belsky, D.W., Higgins-Chen, A., Moskalev, A., Fuellen, G., Cohen, A.A., Bautmans, I., Widschwendter, M., Ding, J., Fleming, A., Mannick, J., Han, J.J., Zhavoronkov, A., Barzilai, N., Kaeberlein, M., Cummings, S., Kennedy, B.K., Ferrucci, L., Horvath, S., Verdin, E., Maier, A.B., Snyder, M.P., Sebastiano, V., Gladyshev, V.N.: Biomarkers of aging for the identification and evaluation of longevity interventions. Cell 186(18), 3758–3775 (2023) 10.1016/j.cell.2023.08.003

[10] Moqri, M., Herzog, C., Poganik, J.R., et al.: Validation of biomarkers of aging. Nat Med 30, 360–372 (2024) 10.1038/s41591-023-02784-9

[11] Mitchell, W., Goeminne, L.J.E., Tyshkovskiy, A., Zhang, S., Chen, J.Y., Paulo, J.A., Pierce, K.A., Choy, A.H., Clish, C.B., Gygi, S.P., Gladyshev, V.N.: Multiomics characterization of partial chemical reprogramming reveals evidence of cell rejuvenation. eLife 12, 90579 (2024) 10.7554/eLife.90579

[12] Tyshkovskiy, A., Kholdina, D., Ying, K., Davitadze, M., Molière, A., Tongu, Y., Kasahara, T., Kats, L.M., Vladimirova, A., Moldakozhayev, A., Liu, H., Zhang, B., Khasanova, U., Moqri, M., Van Raamsdonk, J.M., Harrison, D.E., Strong, R., Abe, T., Dmitriev, S.E., Gladyshev, V.N.: Transcriptomic hallmarks of mortality reveal universal and specific mechanisms of aging, chronic disease, and rejuvenation. bioRxiv (2024) 10.1101/2024.07.04.6019822024.07.04.601982

[13] Holzscheck, N., Falckenhayn, C., Söhle, J., et al.: Modeling transcriptomic age using knowledge-primed artificial neural networks. npj Aging and Mechanisms of Disease 7(1), 15 (2021) 10.1038/s41514-021-00068-5

[14] Martínez-Magaña, J.J., Krystal, J.H., Girgenti, M.J., Núnez-Ríos, D.L., Nagamatsu, S.T., Andrade-Brito, D.E., Group, T.S.B.R., Montalvo-Ortiz, J.L.: Decoding the role of transcriptomic clocks in the human prefrontal cortex. medRxiv (2023) 10.1101/2023.04.19.23288765. Preprint

[15] Meyer, D.H., Schumacher, B.: Bit age: a transcriptome-based aging clock near the theoretical limit of accuracy. Aging Cell 20, 13320 (2021) 10.1111/acel.13320

[16] Buckley, M.T., Sun, E.D., George, B.M., et al.: Cell-type-specific aging clocks to quantify aging and rejuvenation in neurogenic regions of the brain. Nature Aging 3, 121–137 (2023) 10.1038/s43587-022-00335-4

[17] Marioni, J.C., Mason, C.E., Mane, S.M., Stephens, M., Gilad, Y.: Rna-seq: an assessment of technical reproducibility and comparison with gene expression arrays. Genome research 18(9), 1509–1517 (2008)

[18] Wang, L., Feng, Z., Wang, X., Wang, X., Zhang, X.: Degseq: an r package for identifying differentially expressed genes from rna-seq data. Bioinformatics 26(1), 136–138 (2010)

[19] Fang, Z., Cui, X.: Design and validation issues in rna-seq experiments. Briefings in Bioinformatics 12(3), 280–287 (2011)

[20] Love, M.I., Huber, W., Anders, S.: Moderated estimation of fold change and dispersion for rna-seq data with deseq2. Genome biology 15(12), 1–21 (2014) 10.1186/s13059-014-0550-8

[21] Yu, L., Fernandez, S., Brock, G.: Power analysis for rna-seq differential expression studies. BMC Bioinformatics 18(1), 234 (2017) 10.1186/s12859-017-1648-2

[22] Jiang, G., Zheng, J., Ren, S., et al.: A comprehensive workflow for optimizing rna-seq data analysis. BMC Genomics 25, 631 (2024) 10.1186/s12864-024-10414-y

[23] Mboning, L., Rubbi, L., Thompson, M., Bouchard, L.-S., Pellegrini, M.: Bayesage: A maximum likelihood algorithm to predict epigenetic age. Frontiers (2024) 10.3389/fbinf.2024.1329144

[24] Trapp, K.C., Gladyshev, V.N.: Profiling epigenetic age in single cells. Nature Aging 1, 1189–1201 (2021) 10.1038/s43587-021-00134-3

[25] Consortium, T.M.: Single-cell transcriptomics of 20 mouse organs creates a tabula muris. Nature 562(7727), 367–372 (2018) 10.1038/s41586-018-0590-4

[26] Schaum, N., Lehallier, B., Hahn, O., Pálovics, R.e.a.: Ageing hallmarks exhibit organ-specific temporal signatures. Nature 583(7817), 596–602 (2020) 10.1038/s41586-020-2499-2

[27] Gonzalez-Freire, M., Diaz-Ruiz, A., Hauser, D., Martinez-Romero, J., Ferrucci, L., Bernier, M., de Cabo, R.: The road ahead for health and lifespan interventions. Ageing Research Reviews 59, 101037 (2020) 10.1016/j.arr.2020.101037

[28] Culig, L., Sahbaz, B.D., Bohr, V.A.: Effects of lifespan-extending interventions on cognitive healthspan. Expert Reviews in Molecular Medicine 25, 2 (2023) 10.1017/erm.2022.36

[29] Spiridonova, O., Kriukov, D., Nemirovich-Danchenko, N., Peshkin, L.: On standardization of controls in lifespan studies. Aging (Albany NY) 16, 3047–3055 (2024) 10.18632/aging.205604

[30] Han, J.-D.J.: The ticking of aging clocks. Trends in Endocrinology & Metabolism 35(1), 11–22 (2024)

